# Inference of population genetics parameters using discriminator neural networks: an adversarial Monte Carlo approach

**DOI:** 10.1101/2023.04.27.538386

**Authors:** Graham Gower, Pablo Iáñez Picazo, Finn Lindgren, Fernando Racimo

## Abstract

Accurately estimating biological variables of interest, such as parameters of demographic models, is a key problem in evolutionary genetics. Likelihood-based and likelihood-free methods both typically use only limited genetic information, such as carefully chosen summary statistics. Deep convolutional neural networks (CNNs) trained on genotype matrices can incorporate a great deal more information, and have been shown to have high accuracy for inferring parameters such as recombination rates and population sizes, when evaluated using simulations. However these methods are typically framed as regression or classification problems, and it is not straightforward to ensure that the training data adequately model the empirical data on which they are subsequently applied. It has recently been shown that generative adversarial networks (GANs) can be used to iteratively tune parameter values until simulations match a given target dataset. Here, we investigate an adversarial architecture for discriminator-based inference, which iteratively improves the sampling distribution for training the discriminator CNN via Monte Carlo density estimation. We show that this method produces parameter estimates with excellent agreement to simulated data. We developed `dinf`, a modular Python package for discriminator-based inference that incorporates this method, and is available from https://github.com/RacimoLab/dinf/.

## Introduction

In light of the vast quantities of sequencing data being produced today, increasingly complex models are being developed to understand the historical processes that result in the observed genetic data. These population genetic models are necessarily parameterised by unknown quantities, such as past population sizes and migration rates, and many methods have been developed to estimate their values. Evolutionary genetics researchers have the luxury of very sophisticated genetic simulation software that are capable of modelling an enormous variety of biologically-realistic scenarios (Hudson, 2002, Kelleher *et al.*, 2016, Thornton, 2014, Haller & Messer, 2019). While such tools are incredibly valuable, the models being constructed are rarely analytically tractable, so parameter inference can be challenging. One of the most widely-used methods for simulation-based inference is approximate Bayesian computation (ABC) (Beaumont *et al.*, 2002), which has a long history in the field (e.g. see Beaumont, 2010, and references therein). While very effective for some problems, ABC and related methods rely upon comparing summary statistics of the observed genetic data to the simulations, rather than using the full dataset. Furthermore, researchers must first identify statistics that are informative about the parameters being inferred. Inference using analytically tractable models is also possible, but these approaches tend to model only summaries of the genetic data such as the allele frequency spectrum (Gutenkunst *et al.*, 2009, Kamm *et al.*, 2020, Jouganous *et al.*, 2017, Ragsdale & Gravel, 2020, Hernandez & Uricchio, 2015, Excoffier *et al.*, 2021, Steinrücken *et al.*, 2019, Rogers, 2022). Consequently, the vast majority of methods ignore potentially useful information in the observed data.

More recently, it has been shown that deep convolutional neural networks (CNNs) can be used to learn parameters from genotype matrices directly (Chan *et al.*, 2018, Flagel *et al.*, 2019). A genotype matrix is the typical form of the data used to calculate summary statistics, as might be used, for example, in ABC. Such a matrix is high-dimensional and contains essentially all the historically-relevant information in a DNA sequencing dataset. The genotype matrix is also readily produced by genetic simulators, making simulated data an excellent resource for training a CNN. The task is generally framed as a classification problem, or sometimes a regression problem, where the network itself learns which aspects of the data are informative for the parameters in question. Since their introduction to the field, CNNs have been used to infer a variety of biological parameters and evolutionary processes, such as recombination rates, population sizes, migration rates and historical split times, as well as episodes of positive selection and adaptive introgression, to name a few (recently reviewed by Korfmann *et al.*, 2023).

Methods based on neural networks have produced very accurate results, as evaluated using simulations, where the ground truth is known. But many methods rely on well-tuned simulations for training, and model misspecification can be problematic when applied to empirical data (Mo & Siepel, 2023). Moreover, it is difficult to assess how model misspecification impacts the inferences for any specific dataset because, of course, the truth here is not known.

To identify simulation parameters that best fit a dataset, Wang *et al.* (2021) developed a generative adversarial network (GAN; Goodfellow *et al.*, 2014), named `pg-gan`, using a binary classification CNN (the discriminator) with genotype matrices as input. A GAN is a minimax game, in which the output from a data generator is connected to the input of a discriminator—the goal of the discriminator is to distinguish generated data from observed data, and the goal of the generator is to produce data that fools the discriminator. Wang *et al.* (2021) inferred the optimal model parameters for the generator using simulated annealing, which is an iterative optimisation strategy, where in each iteration a new parameter value is proposed and then accepted in a manner similar to a Metropolis-Hastings acceptance criterion. However, `pg-gan` includes an adversarial component in the optimisation procedure, by further training the discriminator CNN whenever a new parameter value is accepted. A GAN approach has also recently been used to detect loci under natural selection. The approach works by training the generator using neutral simulations only, and then finding regions of the genome with low probability of being “fake”, i.e. unlikely to be simulated by the generator (Riley *et al.*, 2023).

To experiment with discriminator-based inference strategies, we developed a modular Python package, named `dinf`. It includes many building blocks needed for constructing and investigating discriminator-based inference methods, such as feature extraction functions, and discriminator CNNs that work well with the extracted features. The source code (https://github.com/RacimoLab/dinf/) is provided under the permissive MIT licence, in the hope that the software, or its components, will be useful to others.

Below, we use Monte Carlo simulation to approximate the output of the discriminator with respect to the model parameters. We show that this straightforward approach produces results comparable to those reported by Wang *et al.* (2021) with `pg-gan`, for a given number of executions of the generator and target functions. However, our approach has the advantage that it explicitly provides an uncertainty estimate such that non-identifiable parameters are more clearly discernible. We then show that the function approximated in one Monte Carlo run can be used as a sampling distribution for the model parameters, which we use to train a more effective discriminator, and subsequently as the sampling distribution for a new Monte Carlo run, in an iterative fashion. This iterative procedure has a strong resemblance to adaptive importance sampling with resampling, and sequential Monte Carlo (SMC; Doucet *et al.*, 2001). Compared with one Monte Carlo run, we find that the iterative approach can produce tighter uncertainty estimates for a given number of executions of the generator and target functions.

## Results

### Discriminator-based inference

We begin by describing the general architecture for discriminator-based inference (Figure 1) and then dive into further details about the specific aspects of our inference framework. A binary-classification neural network (the discriminator) is trained to distinguish between genotype matrices generated via simulation, e.g. with the coalescent simulator `msprime` (Kelleher *et al.*, 2016, Baumdicker *et al.*, 2021), and genotype matrices sampled from a target observed dataset, such as a VCF file (Danecek *et al.*, 2011, Li, 2011) from a genome reference panel, like the 1000 Genomes Project (The 1000 Genomes Project Consortium, 2015). The *n× m* genotype matrices are comprised of *n* haplotypes and *m* loci, where *n* is smaller than the available number of haplotypes in the data, and *m* is small relative to the genome size, such that a large number of distinct genotype matrices may be sampled from the target dataset to train the discriminator. During training, the generated dataset is assigned the class label 0 and the target dataset has the class label 1, so that the trained discriminator predicts the probability that its input *x* comes from the target distribution. Goodfellow *et al.* (2014) show that the output from an optimally trained discriminator is given by

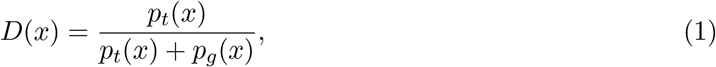

where *p_t_*(*x*) is the density of the target dataset and *p_g_*(*x*) is the density of the generated dataset. To construct the generator dataset for training the discriminator, we first sample simulation parameters *θ* from a distribution, *p_θ_*(*θ*), then the generator is used to produce data with distribution *p_g_*(*x*; *θ*).

**Figure 1:**
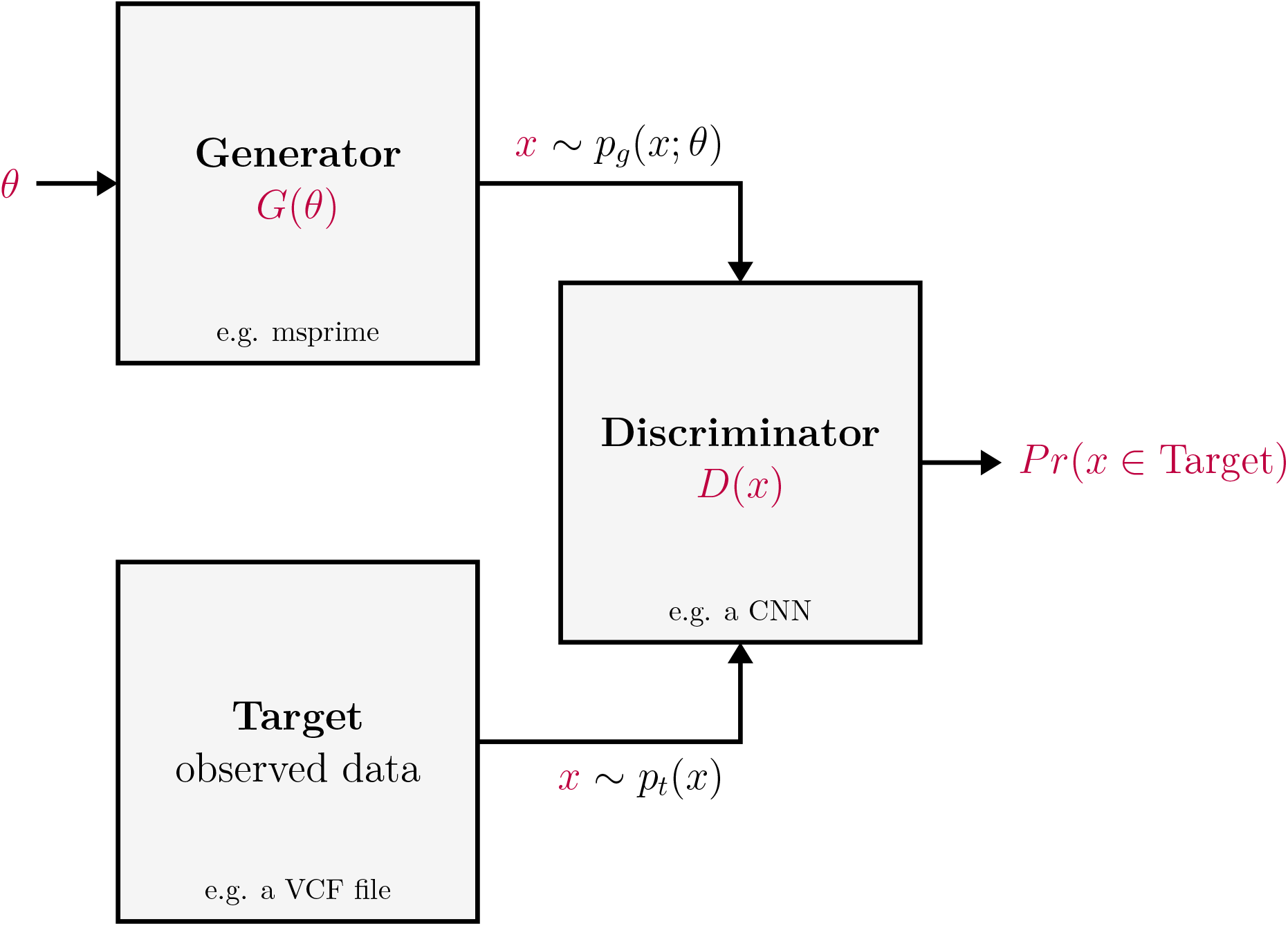
General architecture for discriminator-based inference. A model with *d* parameters, *θ* ∈ℝ^*d*^, is simulated with a black-box generator, *G*, such that *G*(*θ*) produces data from distribution *p_g_*(*x*; *θ*). Similarly, data with distribution *p_t_*(*x*), is sampled from a target dataset (typically observed data). The discriminator, *D*, is a binary classifier that is trained to distinguish the generated data distribution, *p_g_*(*x*; *θ*), from the target data distribution, *p_t_*(*x*). The goal of inference is to find values *θ*^*^ such that the discriminator is not able to distinguish *p_g_*(*x*; *θ*^*^) from *p_t_*(*x*).

The distribution of the generated data used in training is

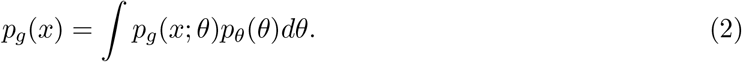

Shifting focus to the generator, we wish to generate data using some optimal set of parameters *θ*^*^, such that the trained discriminator cannot distinguish the generated dataset from the target dataset. One way to find *θ*^*^ is to maximise the objective function,

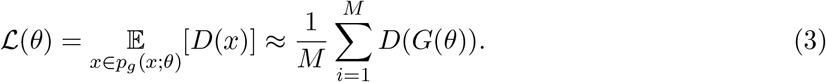

In the GAN described by Goodfellow *et al.* (2014), the generator function, *G*, is differentiable, so the analogous objective was optimised using stochastic gradient descent. However, our generator may be an external program and must be treated as a black box. Thus gradients are not available in closed form, nor using automatic differentiation techniques. In `pg-gan`, Wang *et al.* (2021) maximise this objective using simulated annealing, because it doesn’t rely on the calculation of gradients.

For GANs where the generator is also a neural network, *θ* represents the trainable network weights and is very high dimensional (e.g. more than 1 million parameters), so training the discriminator by sampling from *p_θ_*(*θ*) in Equation (2) is not practical. Instead, Goodfellow *et al.* (2014) alternate between training the discriminator using a point mass for *p_θ_*(*θ*), such that *p_g_*(*x*) = *p_g_*(*x*; *θ*^(*i*)^) for some value *θ*^(*i*)^, then doing one step of optimising parameters for the generator via Equation (3). In effect, the generator uses the discriminator output to improve the data generation process, by proposing new parameter values, *θ*^(*i*+1)^, that are presumably closer to *θ*^*^. Then in the next iteration, the discriminator is trained with data from the improved generator, 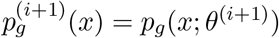, thereby increasing discriminator accuracy in this region of parameter space. Following an initial round of discriminator training, `pg-gan` uses this same approach of training the discriminator based on the current best guess for parameters, *θ*^(*i*)^. However, if *θ*^(*i*)^ is not very close to the optimal *θ*^*^, the discriminator may still have limited accuracy in regions near *θ*^*^, so convergence may be slow. For improved convergence, we would like for *p_θ_*(*θ*^*^) to always be non-zero when training the discriminator, and ideally *θ*^*^ = arg max *p_θ_*(*θ*). With this in mind, we construct a sequence of non-point-mass sampling distributions 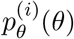, that converge towards a point mass at *θ*^*^. We first approximate the discriminator output as a function of *θ* using Monte Carlo simulation and kernel density estimation (KDE). The approximation is then used to define the sampling distribution, 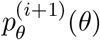, for training the discriminator in the next iteration. This focuses discriminator training on the region of parameter space which is most likely to contain the optimal parameter *θ*^*^, by explicitly sampling in proportion to the discriminator output. Pseudocode for this procedure is provided in Algorithm 1. To train the discriminator in the first iteration, set the initial sampling distribution to *π*(*θ*), a starting distribution with *π*(*θ*^*^) > 0,

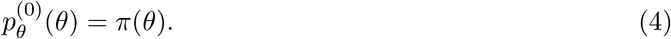

For iteration *i*, we have

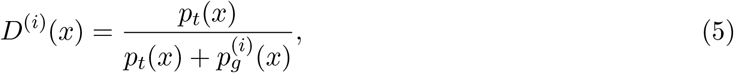

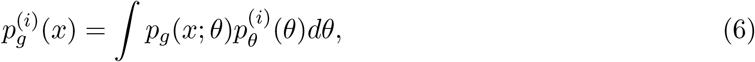

The sampling distribution for the next iteration is then given by

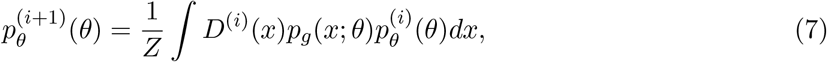

where *Z* is chosen to make the integral sum to 1. We approximate the sampling distribution in Equation (7) using a KDE with kernel *K*, of samples drawn from 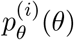, weighted by the discriminator prediction *D*^(*i*)^(*G*(*θ*)),

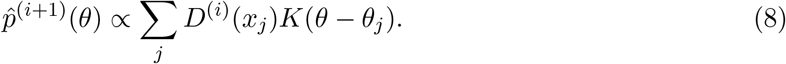

In practice, the *d*-dimensional KDE is not computed explicitly in each iteration, because the multidimensional KDE is only used for sampling. Instead, samples are drawn with replacement from the previous iteration’s proposals in proportion to the discriminator predictions, and smoothed according to the KDE kernel function (i.e. by adding *d*-dimensional Gaussian noise). For visualising the sampling distributions, 1-dimensional marginal KDEs are computed explicitly.

#### Algorithm 1

Adversarial Monte Carlo sampler. For clarity of exposition, we assume that the functions *G*(*θ*) and *D*^(*i*)^(*x*) accept and produce vectors.

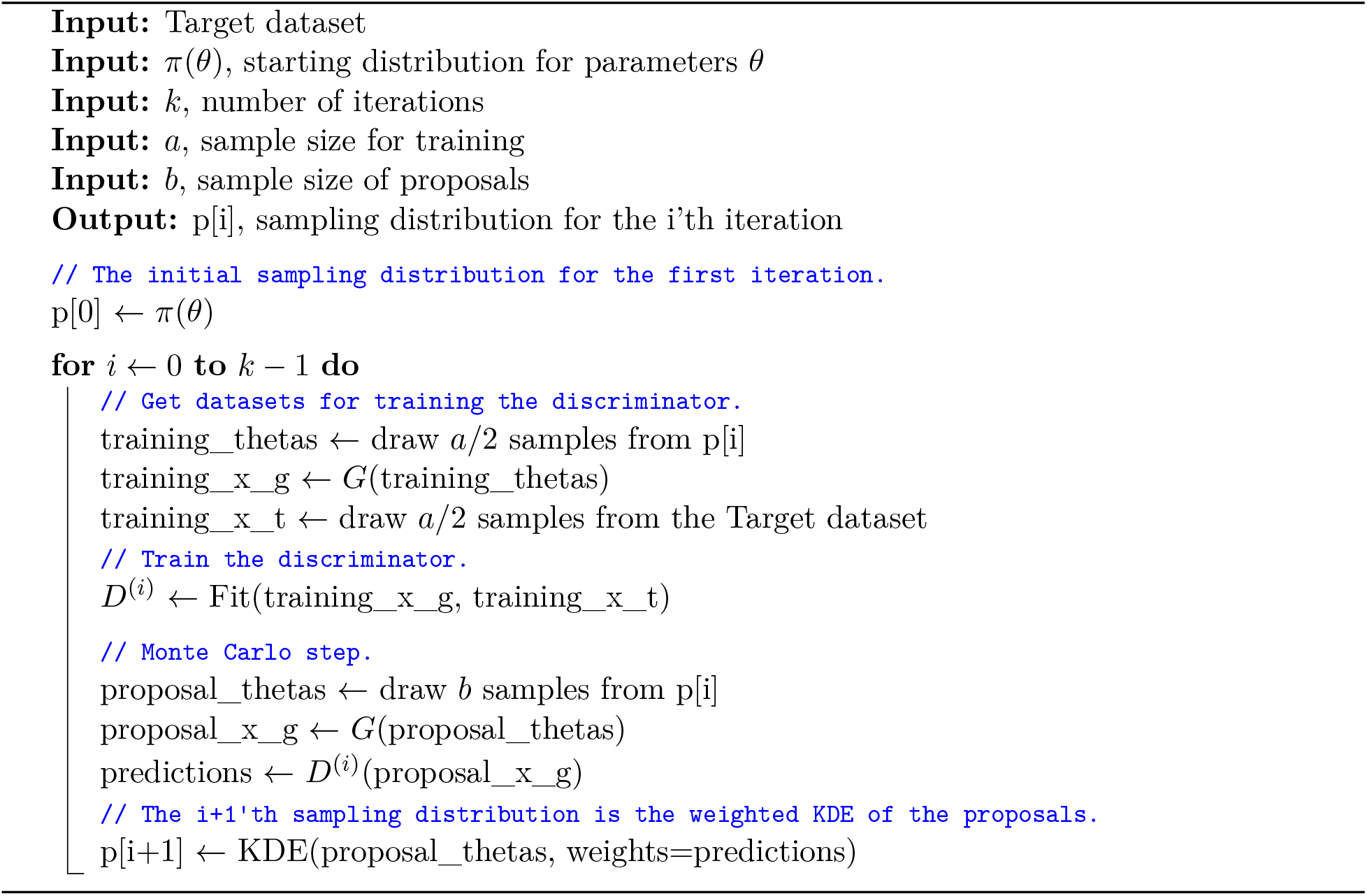

### Application to an isolation-with-migration model

We implemented the six-parameter isolation-with-migration model (Figure S1 and Table S1) that was used to evaluate `pg-gan` (Wang *et al.*, 2021). The inferrable parameters in this model include the recombination rate (reco), the population sizes in the ancestral and two extant populations (N_anc, N1, N2), the split time between the two present-day populations (T_split), and the proportion of migrating individuals for a pulse of migration between the two present-day populations (mig). We used a uniform prior on each parameter, with upper and lower bounds matching those used by Wang *et al.* (2021). We described the demographic model using a templated demes (Gower *et al.*, 2022) model, which was passed to msprime (Kelleher *et al.*, 2016, Baumdicker *et al.*, 2021) for simulation. For the discriminator, we employed a permutation-invariant convolutional neural network (CNN) that treats each haplotype within a population as exchangeable (Chan *et al.*, 2018). To assess the performance of the discriminator, we sampled from the target dataset by calling the generator function with a fixed set of values, *θ*^*^, the “truth” values. Through experimentation with this model and others, we converged on a shallow CNN architecture (see Methods) with ∼30,000 trainable parameters per sampled population (the two present-day populations, for the isolation-with-migration model).

### Monte Carlo density estimation

To assess the plausibility of doing Monte Carlo density estimation with Equation (8), we first evaluated the results from a single density estimation step (i.e. without iteration) when using a large number of samples. We trained a discriminator for 20 epochs on 1 million replicates: 500,000 from the target dataset and 500,000 from the generator, where parameter values for the generated data were drawn from uniform prior distributions (Table S1). The discriminator achieved a training accuracy of 99.9%. Evaluation on a test dataset of 100,000 instances (50,000 from the target dataset, 50,000 from the generator) produced a test accuracy of 99.7%, and showed negligible signs of overfitting (Figure S2a). We then drew 1 million fresh samples from the generator and made predictions with the discriminator. The distribution of discriminator predictions (Figure S2b) was strongly multi-modal, with most samples having a prediction near zero, and a smaller peak of predictions near one. In total, this took 8.5 hours to run with a peak RAM use of 21.3 Gb on an 80 core Xeon 6248 server with a Tesla T4 GPU for training the discriminator and making discriminator predictions.

Finally, we obtained a marginal distribution of Equation (8) for each parameter by constructing weighted histograms and weighted KDEs (Figure 2). The recombination rate and the ancestral population size were estimated very effectively, however the migration parameter was estimated poorly. Distributions for the split time and population sizes of the extant populations were centred on the “truth” values, although the median values were not as close to the truth as for the recombination rate and the ancestral population size parameters.

**Figure 2:**
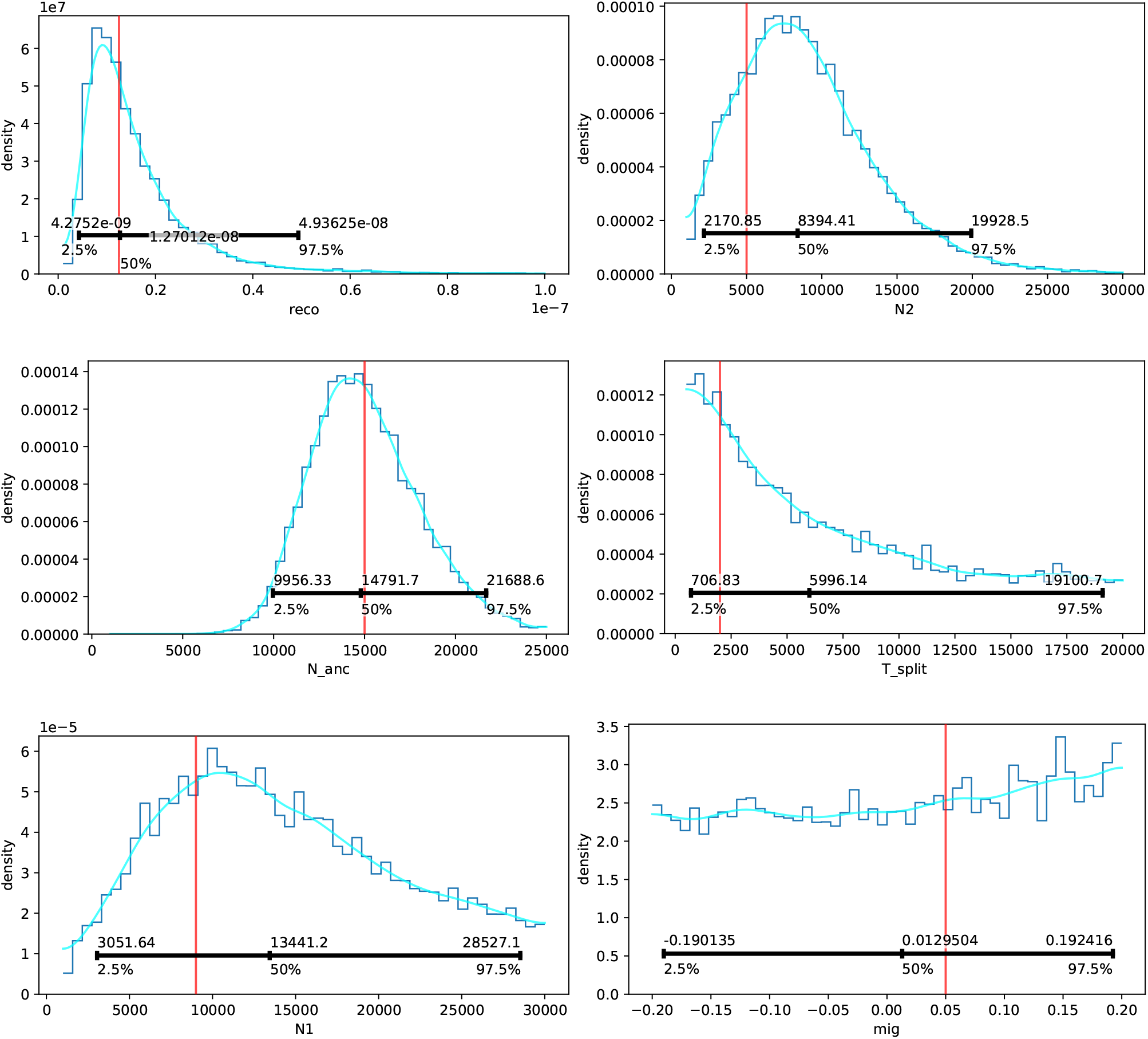
Marginal distributions of the discriminator output for each of the six inferrable parameters of an isolation-with-migration model, approximated via Monte Carlo density estimation. In each panel: the dark blue line shows the marginal histogram; the light blue line shows the marginal KDE; the red vertical line indicates the “truth” value; and the horizontal black bar with whiskers shows the median and 95% compatibility intervals of the sample.

### Adversarial Monte Carlo

Encouraged by the one-step Monte Carlo results, we performed inference on the same isolation-with-migration model for 30 iterations, with fewer samples per iteration. The discriminator was trained for five epochs in each iteration, on a training dataset of 100,000 replicates. The sampling distributions were constructed from a separate proposal dataset of 100,000 replicates in each iteration, using a Gaussian KDE with bandwidth chosen using Scott’s rule of thumb (Scott, 2015), and kernel shrinkage according to West (1993). Thus, 10 iterations of the adversarial Monte Carlo analysis corresponds to 1 million replicates for training and 1 million replicates for proposals, which is equivalent to that used for the one-step Monte Carlo density estimate (Figure 2). In total, this took 6 hours 15 minutes to run with a peak RAM use of 11.4 Gb on an 80 core Xeon 6248 server with a Tesla T4 GPU for training the discriminator and making discriminator predictions.

We constructed a test dataset of 100,000 replicates, and evaluated the discriminator against this test dataset in each iteration. Training metrics (Figure S3) indicated a high training accuracy of 99.3% in the first iteration, which progressively declined in each iteration, appearing to converge to 68.5% in the final iteration. The test accuracy was also 99.3% in the first iteration, and test accuracy declined to a final test accuracy of 83.6%, although the test metrics were a lot more variable than the training metrics.

We created a violin plot of the sampling distributions in each iteration (Figure 3). After around 10 iterations, we saw excellent agreement between the distribution medians and the truth values (indicated in red) for all parameters except the migration rate. In further iterations, the sampling distributions became narrower, and remained centred on the truth values (except for the migration rate). All distributions for the migration rate stubbornly remained similar to the uniform starting distribution.

**Figure 3:**
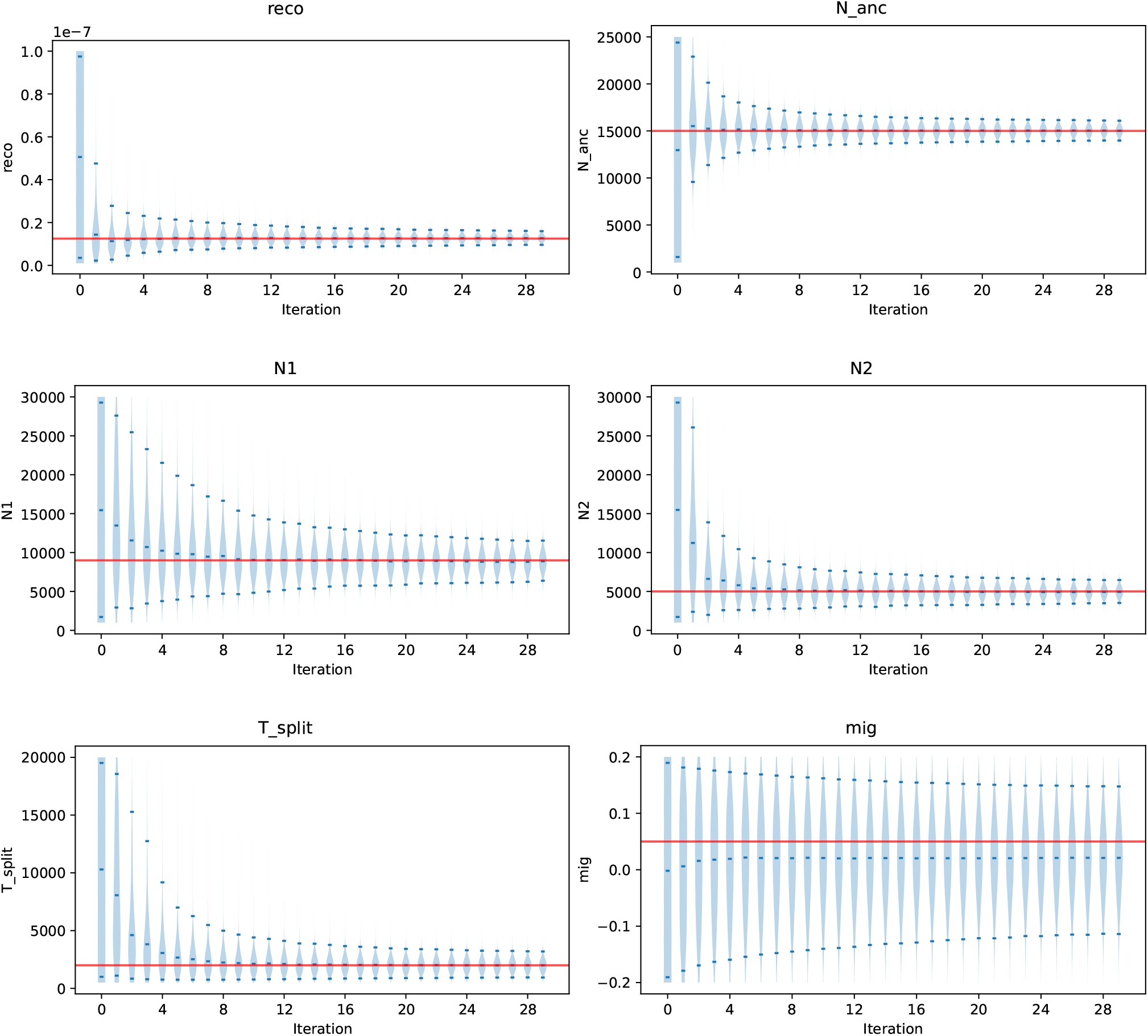
Adversarial Monte Carlo sampling distributions for the six inferrable parameters of an isolation-with-migration model. In each panel: the violin plot shows a marginal KDE for each iteration (light blue); the red horizontal line indicates the “truth” value; dark blue marks within each KDE show the median and 95% compatibility intervals.

To assess convergence of the sampling distributions, we calculated the entropy relative to uniformity for the discriminator predictions obtained in each iteration (Figure S4), as suggested by West (1993). When the discriminator predictions provide no additional information compared with the sampling distribution, the relative entropy should approach a value of 1.0. For the isolation with migration model, we see that the relative entropy is quite close to 1.0 from around iteration 10 onward. While our interpretation is that the sampling distribution has converged, we caution that this does not provide evidence that the distribution has converged to any specific distribution.

### Simulated annealing

We initially set out to improve sampling efficiency via an iterated MCMC, rather than via Monte Carlo density estimation, using the discriminator output (Equation (3)) to define a Metropolis-Hastings acceptance condition. However, this suffered from convergence issues (results not shown). We note that `pg-gan’s` optimisation via simulated annealing has a strong resemblance to MCMC, in that it uses a very similar criterion to choose whether to accept a proposal. To better understand `pg-gan`, and the impressive results reported in Wang *et al.* (2021), we reimplemented their algorithm within our inference software. However, we were unable to reliably infer the migration rate parameter.

To confirm that this was not an issue with our reimplementation, we did 10 runs of `pg-gan` on the isolation-with-migration model, and calculated the absolute error of the parameter estimates, divided by the expected error from guessing uniformly at random (Figure S5a). By normalising according to the expected error from guessing, we are able to more fairly compare estimates between parameters with different scales for their prior distributions. From Figure S5a, we see that the migration rate estimates are typically the worst estimates of all parameters—only slightly better than guessing. In each iteration, `pg-gan` constructs 10 distinct proposals for each parameter, while leaving the remaining parameter values fixed, then chooses the best proposal among all parameters by ranking them according to the discriminator output (Equation (3)). We found that proposals which modified the migration rate parameter were the least frequently accepted of all parameters (Figure S5b), suggesting that this parameter has relatively little influence on the discriminator output.

In each iteration, `pg-gan` drew 3000 samples from the generator for proposals (*M* = 50 replicates for each of 10 proposals for each of 6 parameters). When a proposal is accepted, the discriminator was trained on a further 5000 samples from each of the generator and the target. In our 10 runs, we observed an average acceptance rate of 0.6 (per iteration). Additionally, the discriminator was pre-trained on 50,000 samples from each of the generator and target before iterating. So for 300 iterations (the `pg-gan` default), this corresponds to a total of 1.85 million calls to the generator and 950,000 calls to the target function—more than that used for the one-step Monte Carlo analysis above.

### Dinf software package

We developed a modular Python package named dinf, for discriminator-based inference, with a Python interface for defining models, and a command line interface for performing inference and plotting results. Users can write models by defining the parameters to be inferred, writing a generator function, and writing a target function. Models can include one or multiple sampled populations. Functions are provided to build genotype matrices from tskit (Kelleher *et al.*, 2018) succinct tree sequences, as supported by msprime (Kelleher *et al.*, 2016, Baumdicker *et al.*, 2021), fwdpy11 (Thornton, 2014) and SLiM (Haller *et al.*, 2019) simulators. Genotype matrices can also be sampled directly from VCF or BCF files (Danecek *et al.*, 2011, Li, 2011), with no need for data conversion a priori. Calls to the generator and target functions execute in parallel, to take advantage of many-core systems. The discriminator neural networks are implemented using JAX (Bradbury *et al.*, 2018) and Flax (Heek *et al.*, 2020), and users may customise the provided networks or write their own from scratch. For inference, dinf implements iterated Monte Carlo, MCMC, and `pg-gan-style` simulated annealing. Dinf is installable from the pypi and conda-forge repositories and the source code at https://github.com/RacimoLab/dinf/ is distributed under the permissive MIT licence.

## Discussion

### Adversarial Monte Carlo

We found that iteratively applying Monte Carlo density estimation is very effective for demographic parameter inference via discriminator functions. Not only does it provide accurate estimates of the parameter values and explicit uncertainty estimates, it also allows for more effective training of the discriminator via importance sampling of the training dataset in each subsequent iteration. As the iterations proceed, the sampling distribution focuses in on a small region of parameter space, where the discriminator is least able to distinguish generated data from the target dataset. Once the discriminator can no longer distinguish the two classes of data, then *D*^(*i*)^(*G*(*θ*)) = 0.5 for all 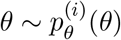, so the sampling distribution will no longer change. This makes our proposed adversarial procedure very stable, in comparison to training a regular GAN.

As a corollary, if the data contains no information about a given parameter, such that the discriminator cannot learn anything about that parameter, the sampling distributions will simply resemble the starting distribution. The latter was observed for inferences of the migration rate parameter (Figures 2 and 3), which suggests it isn’t readily identifiable from the genomic data provided to the discriminator. One way to improve inference of this parameter could be to model an additional sister group to deme2 that is not the recipient of gene flow from deme1.

For the KDE bandwidth, we used Scott’s rule of thumb, which is appealing due to its ease of application, but it produces density estimates that are oversmoothed, particularly in high dimensions. This may explain why the discriminator is still able to distinguish generated data from the target dataset in later iterations (see training loss and accuracy in Figure S3), in spite of the appearance of convergence (Figure S4). Alternatives, such as an adaptive bandwidth, which has different values in different regions of parameter space, may be worth exploring to further improve convergence. We also note two additional technical choices that affect the resulting density approximation: (1) the sampling distributions were truncated to upper and lower bounds (Table S1) by resampling values that were out of bounds for any parameter, and (2) we applied kernel shrinkage as described by West (1993) to avoid inflation of sampling variance in each iteration. A consequence of (1) is that the density is reduced near parameter bounds, and (2) works by “shrinking” sample values towards the distribution mean. For the estimation of the migration rate parameter, it is possible that one or both of these choices have artificially narrowed the sampling distributions compared to the uniform starting distribution, rather than this being the result of the discriminator predictions themselves.

### Simulated annealing and pg-gan

To choose whether to accept a proposed *θ*^(*i*^*′*^)^, `pg-gan` compares the discriminator output of the proposal to that of the current position *θ*^(*i*)^. For this to work, a reasonably accurate approximation to the objective function (Equation (3)) is needed, which `pg-gan` calculates by taking an average over a moderate number of draws (50) from *G*(·). Thus this method requires a substantial number of calls to the generator over a realistic number of GAN iterations.

Training the GAN can be tricky, as it requires balancing the interests of both the generator and the discriminator so that neither dominates the other. To this end, the `pg-gan` discriminator architecture was intentionally not optimised. For example, the sum function was chosen for the permutation-invariant layer, which was known to slow down training when compared to the max function.

In each iteration where a proposed *θ*^(*i*^*′*^)^ is accepted, `pg-gan` trains the discriminator further on samples from *p_g_*(*x*; *θ*^(*i*^*′*^)^). As an unintended consequence, the discriminator may now produce poorer inferences when applied to values farther from *θ*^(*i*^*′*^)^. Hence the values of the objective function that are calculated for proposals in early iterations, when the temperature parameter is still high, can potentially be erratic. As the temperature parameter is decreased in every iteration, proposals are increasingly closer to the current position, so this issue should diminish as the algorithm proceeds. An alternative strategy that may ameliorate this effect could be to instead train the discriminator on parameter values sampled from a ball around *θ*^(*i*^*′*^)^.

### Limitations

One way of viewing the role of the discriminator, is that it performs dimensionality reduction on the genetic data to produce a single informative summary statistic. In this sense, the discriminator is very effective at mitigating the curse of dimensionality of the summary statistics, which is problematic for ABC. On the other hand, the discriminator has no impact on the dimensionality of the space of model parameters. While we were able to reasonably perform inference for a six-parameter model, convergence speed may yet make the method impractical for very complex models with several tens of parameters.

Model misspecification remains a potential issue with discriminator-based inference. In particular, we recognise that data missingness, genotype error, SNP ascertainment, unmodelled population structure, and/or incomplete model specification (e.g. neutral-only mutations) will all contribute to differences between empirical datasets and simulated datasets (Johri *et al.*, 2022a). While many of these differences should be discernible by the discriminator, inference will still be possible in many cases, albeit with possibly reduced rates of convergence. Furthermore, we hope that discriminator-based inference can be used as an aid to developing simulation models for data missingness, genotyping error and SNP ascertainment.

Discriminator-based inference relies on *sampling* from the target dataset to train the discriminator. This means that each input datum is necessarily limited to a subset of the entire dataset, which for our purposes is a (short) window of the genome. This does limit the information that can be automatically extracted from the dataset, such as information about long tracts of identity-by-state or long range linkage disequilibrium. While long range correlations could be observed by using gapped genotype matrices (i.e. multiple genomic windows, with a fixed-size gap between them) or other creative solutions, this would need careful consideration of how the parameters in question might influence the data.

### Future prospects

Many processes that biologists wish to explore cannot be adequately modelled in a coalescent framework, e.g. distributions of fitness effects (Johri *et al.*, 2022b, Matheson & Masel, 2023) or complex geographical movement (Battey *et al.*, 2020, Petr *et al.*, 2023). These processes can be modelled with forwards-in-time simulators (Thornton, 2014, Haller *et al.*, 2019), but at the expense of vastly increased simulation time compared to a coalescent simulator. Thus for many biologically relevant models, simulation time is expected to dominate the computational cost. To reduce the number of simulations required for parameter inference, there are multiple avenues for future improvements.

There have been many recent advances in simulation-based inference, particularly focused on conditional density estimation using neural networks (recently reviewed by Cranmer *et al.*, 2020). With neural density estimation, the network is trained on simulations to produce an estimate for the parameters of a parameterised density, which approximates the likelihood, or alternately the posterior, of *θ*. In contrast, density estimation in our discriminator-based inference framework is done outside of the network using a kernel density estimate, and the density we infer is not interpretable as a likelihood or a posterior. Neural density estimation also shares similarities with networks framed as a regression task (Sanchez *et al.*, 2023), in that the observed data is not incorporated during training. These networks are referred to as “armortized”, meaning that once trained they can be subsequently applied to many distinct observations in a dataset, rather than being focused on a specific observation. For genomic datasets, this means that the network could be trained once and then applied multiple times on different regions of the genome. On the other hand, because the inference is not focused, the results may not be well calibrated for the parameter space corresponding to any one observed datum. In terms of both the number of simulations required for inference, and accuracy, neural density estimation approaches compare favourably to more traditional ABC and ABC-SMC approaches (Lueckmann *et al.*, 2021). Neural density estimation can also be used in a iterative manner to focus inference on a specific observation, by using the posterior density for a given observation as the prior in a subsequent iteration. As we observed here for Monte Carlo, iterative approaches (including ABC-SMC and iterated neural density estimation) tend to produce superior results for a given simulation budget, when compared to their non-iterative counterparts (Lueckmann *et al.*, 2021).

A variety of encodings for genotype data and network architectures have been successfully applied in population genetics, so it seems likely that the exact choices here are somewhat arbitrary. But for any simulation-based inference method using a neural network, the choice of encoding and corresponding network architecture will undoubtedly influence the number of simulations required to obtain satisfactory parameter estimates and measures of uncertainty. For inference methodology to progress with greater confidence, it will be essential to first assess if the genotype matrix (or any specific encoding thereof) constitutes a sufficient statistic for the parameters in question. Assessing this in the general case is within our grasp (e.g. see Chen *et al.*, 2021), and such information-theoretic investigations can also inform users about the appropriate size of the genotype data needed for a given problem (i.e, the number of individuals or number of loci to simulate) (Kim *et al.*, 2019). Likewise, the network architectures themselves should be assessed for sufficiency. Exchangeable CNNs (Chan *et al.*, 2018) provide a very reasonable inductive bias (Battaglia *et al.*, 2018) for extracting information from genotype matrices, but so too might other architectures. For instance, transformer architectures (Vaswani *et al.*, 2017) are largely unexplored in population genetics, but in the wider machine learning literature transformers have essentially replaced recurrent networks for natural language problems, and also shown state-of-the-art performance on computer vision problems where CNNs have historically dominated (Dosovitskiy *et al.*, 2023).

While we have not developed any strategies for model comparison here, several techniques should be straightforward to apply. A simple goodness of fit measure is the average discriminator prediction for samples drawn from the target dataset. As better models should produce greater discriminator confusion, a lower prediction value would indicate a superior model. This would need care in application, as it assumes an equivalent amount of discriminator training for each model. An arguably more robust approach could be to do a reciprocal comparison using generated data, by giving data generated under model A to the discriminator for model B, and vice versa. Another approach would be to include a categorical variable as an inferrable parameter that chooses between a set of competing models (Toni *et al.*, 2009, Toni & Stumpf, 2010). We can also imagine doing automatic exploration of model space in the spirit of GADMA (Noskova *et al.*, 2020, Noskova & Borovitskiy, 2022).

Overall, our adversarial Monte Carlo approach for discriminator-based inference performs very well at demographic inference, but also leaves open several avenues for further expansion and improvement. Its associated modular Python package—dinf—can be used to deploy this approach on other sets of genomic data, as well as for constructing new discriminator-based inference frameworks using easy-to-use modular code. Adversarial methods in general appear to be quite powerful at extracting historical information from genotype data, and illustrate that we are yet to fully grasp the scope of what CNNs can achieve when deployed for evolutionary genetic inference.

## Methods

### Simulations

The isolation-with-migration model (Figure S1 and Table S1) was simulated with msprime v1.2.0 (Kelleher *et al.*, 2016, Baumdicker *et al.*, 2021), using a demes (Gower *et al.*, 2022) YAML string for the demographic model, and substituting parameter values into the string with Python’s builtin templating support (Van Rossum & Drake Jr, 2009). We simulated a 1 Mbp region using a mutation rate of 1.25e-8 per base pair per generation, and sampled 32 diploid individuals from each of the model’s two present-day populations. We treated the data as phased and did not apply any filtering for low-frequency alleles. A genotype matrix was extracted for each sampled population in the simulation output. Each matrix was constructed as a binned haplotype matrix with 128 bins, following Gower *et al.* (2021). Briefly, we built an *n × m* matrix, by partitioning the *n* haplotypes into *m* bins of equal length, and counting minor alleles in each bin. Compared with using a fixed number of SNP sites, this avoids the need to separately encode inter-SNP distances, and more compactly encodes a larger genomic region, at the expense of decreased fine-scale fidelity.

### Discriminator network

For the discriminator, we used an exchangeable convolutional neural network with multiple inputs— one genotype matrix for each of the sampled populations. These matrices were processed separately, and combined only in the final output layer of the network. Each matrix in an input batch was a multidimensional array, with dimensions (*batch, haplotype, loci, channels*), where the *haplotype* and *loci* dimensions correspond to the *n* × *m* dimensions of the genotype matrix, and the final dimension nominally has size 1. Two 1-dimensional convolution layers with 32 and 64 features respectively were successively applied along the *haplotype* dimension and feature size was reduced in each convolution using a stride of 2 units. Then the max function was used to collapse information across the *haplotype* dimension, making the network invariant to the order of haplotypes (Chan *et al.*, 2018). A third 1-dimensional convolution followed, this time operating on the *loci* dimension (with 64 features, and stride of 1 unit), and the max function was then used to collapse information across the *loci* dimension. Each convolution kernel had a size of 5 units. ELU (exponential linear unit) activation was used after each convolution layer, and the final output which was transformed into a probability on [0, 1] with the sigmoid function. The number of trainable parameters in the network was dependent on the number of matrices in the input (i.e. the number of populations), but independent of the size of the genotype matrices, for a total of 31,589 per input matrix. We included a batch normalisation layer prior to each convolution, which served to normalise the input data, to improve the robustness of the network with respect to possible gradient explosion, and to mitigate over-fitting. The network was trained using the Adam optimiser with learning rate 0.001, binary cross entropy for the loss function, and gradients averaged over input batches of size 64.

### pg-gan

We ran pg-gan (git commit b1a82ae) 10 times, with seeds=1,…,10 for the random number generator.

## Acknowledgements

We thank Rasa Muktupavela and Sara Mathieson for helpful comments on the text. GG was funded by the European Commission in the framework of the Horizon2020 Project FindingPheno (GA 952914). FR was supported by a Villum Young Investigator Grant (project no. 00025300), a COREX ERC Synergy grant (ID 951385) and a Novo Nordisk Fonden Data Science Ascending Investigator Award (NNF22OC0076816).

## Supplementary material

**Figure S1:**
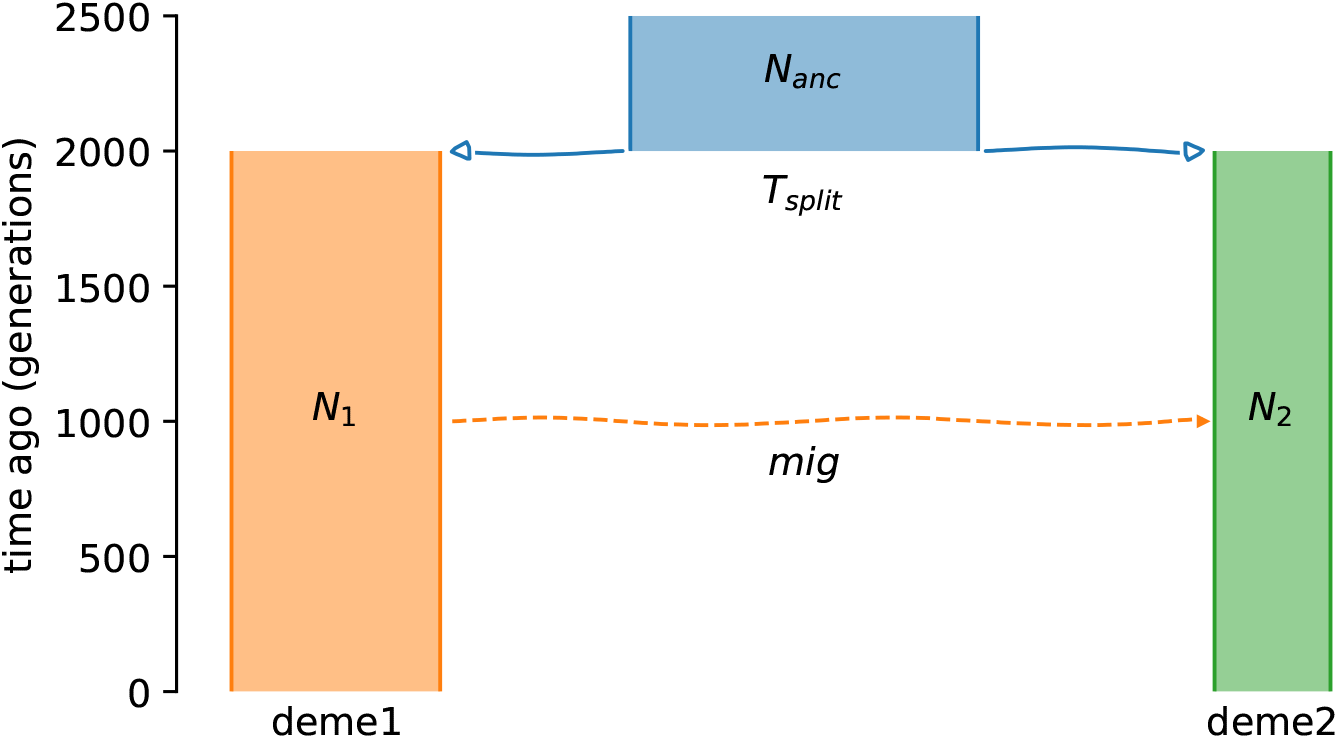
A six-parameter isolation-with-migration demographic model, as used by Wang *et al.* (2021) to evaluate `pg-gan`. The time of the migration pulse is not a free parameter—it is fixed at time T_split/2.

**Table S1:** Free parameters for the isolation-with-migration demographic model.

| Parameter | Truth | Prior | Description |
| --- | --- | --- | --- |
| **reco** | 1.25e-8 | Unif(1e-9, 1e-7) | Recombination rate |
| **N_anc** | 15000 | Unif(1000, 25000) | Ancestral population size |
| **N1** | 9000 | Unif(1000, 30000) | Population size of deme1 |
| **N2** | 5000 | Unif(1000, 30000) | Population size of deme2 |
| **T_split** | 2000 | Unif(500, 20000) | Split time for deme1 and deme2 |
| **mig** | 0.05 | Unif(-0.2, 0.2) | Proportion of migrating individuals between deme1 and deme2 at time $T_{split}/2$. Positive values indicate migration from deme1 to deme2, negative values from deme2 to deme1. |

**Figure S2:**
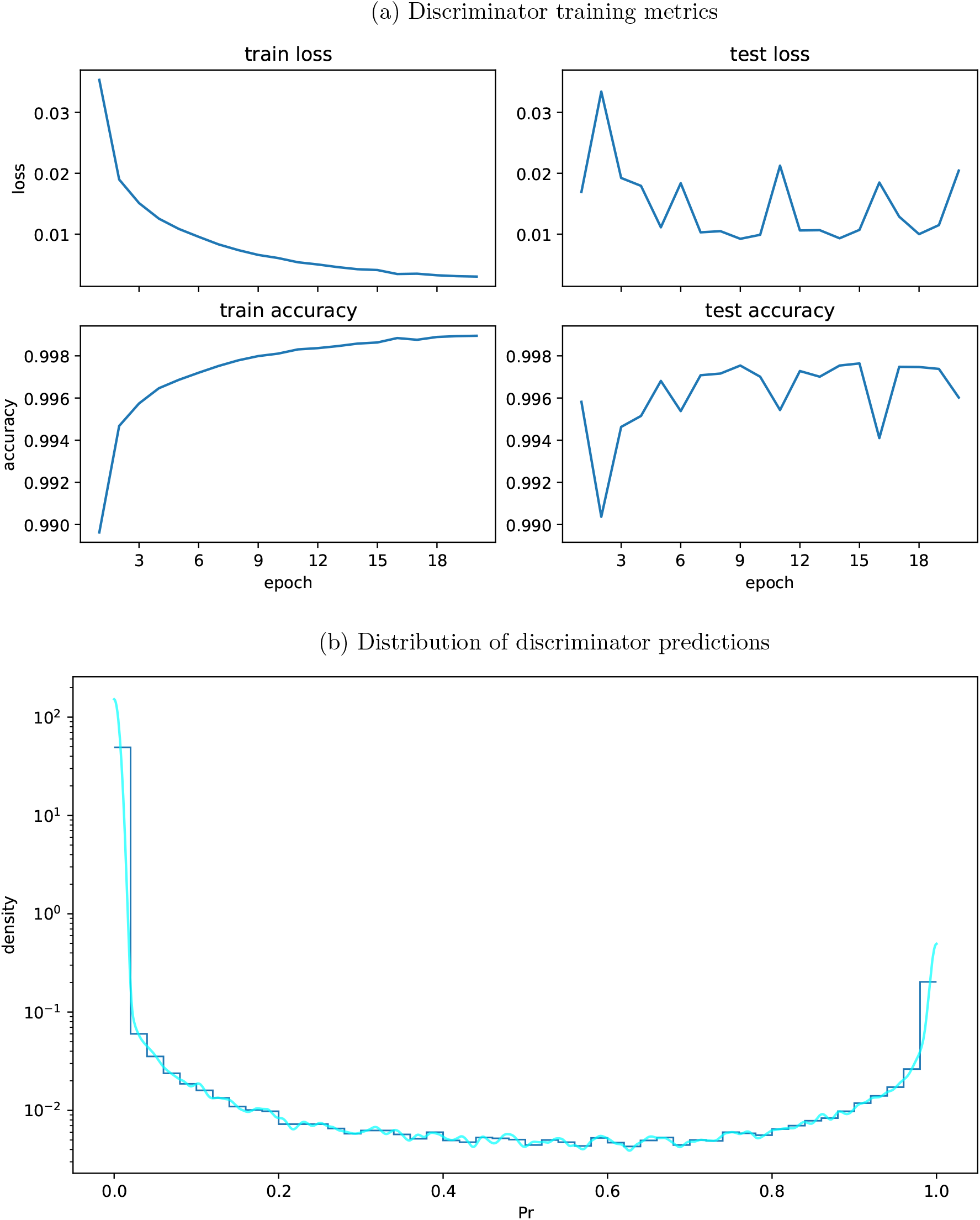
Monte Carlo simulation of isolation-with-migration model. (a) Discriminator training metrics for a training dataset of 1 million replicates (500,000 from the target dataset, 500,000 from the generator) and a test dataset of 100,000 instances (50,000 from the target dataset, 50,000 from the generator). (b) Distribution of discriminator predictions on a dataset of 1 million proposals from the generator. Dark blue shows a histogram. Light blue shows a KDE.

**Figure S3:**
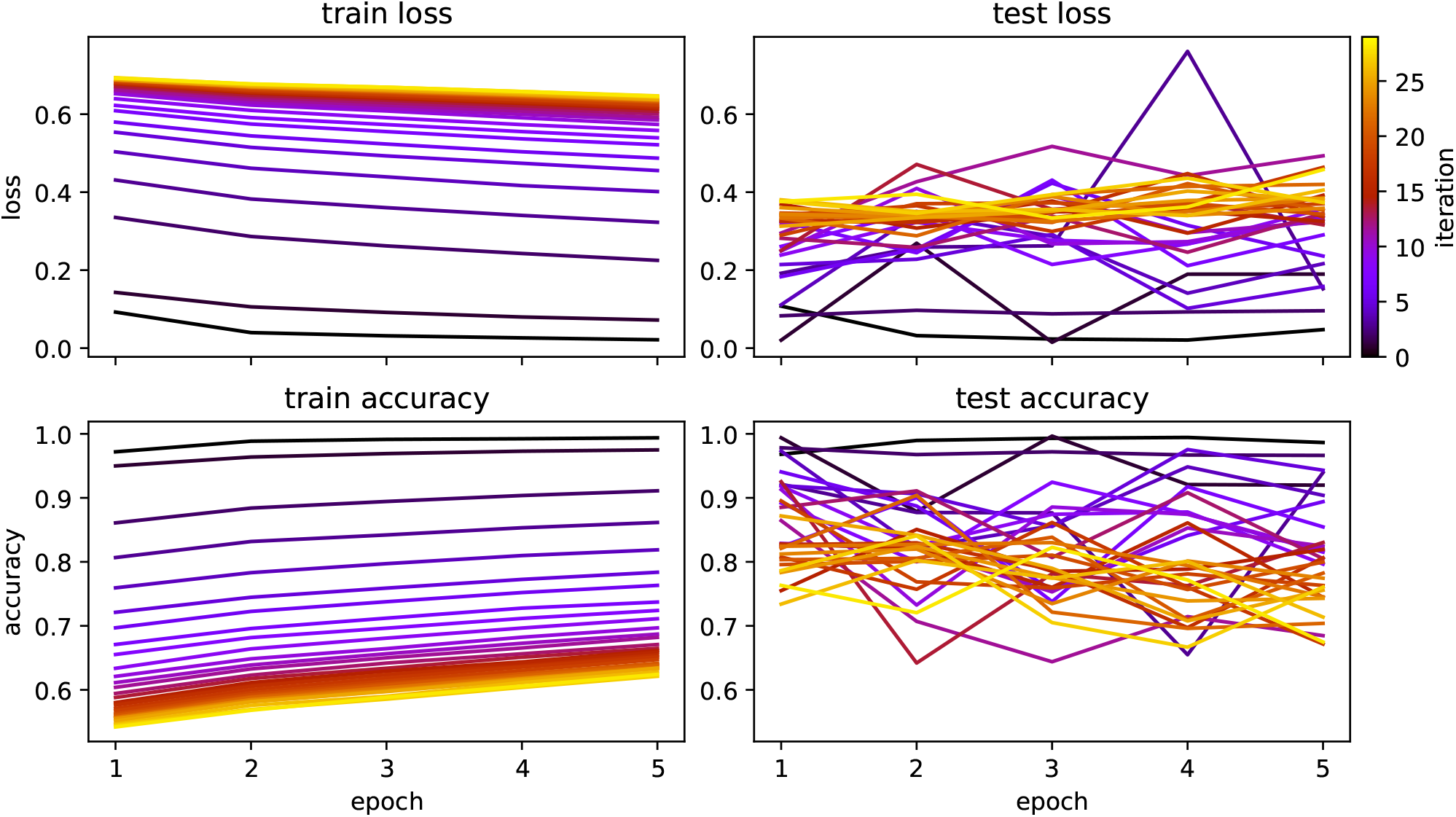
Training metrics for discriminator networks trained on the isolation-with-migration model in each adversarial Monte Carlo iteration. The discriminator was (further) trained on 100,000 replicates (50,000 from the target dataset, 50,000 from the generator) in each iteration. A test dataset of 100,000 replicates (50,000 from the target dataset, 50,000 from the generator) was constructed up front, and the discriminator was evaluated on this test dataset in each iteration.

**Figure S4:**
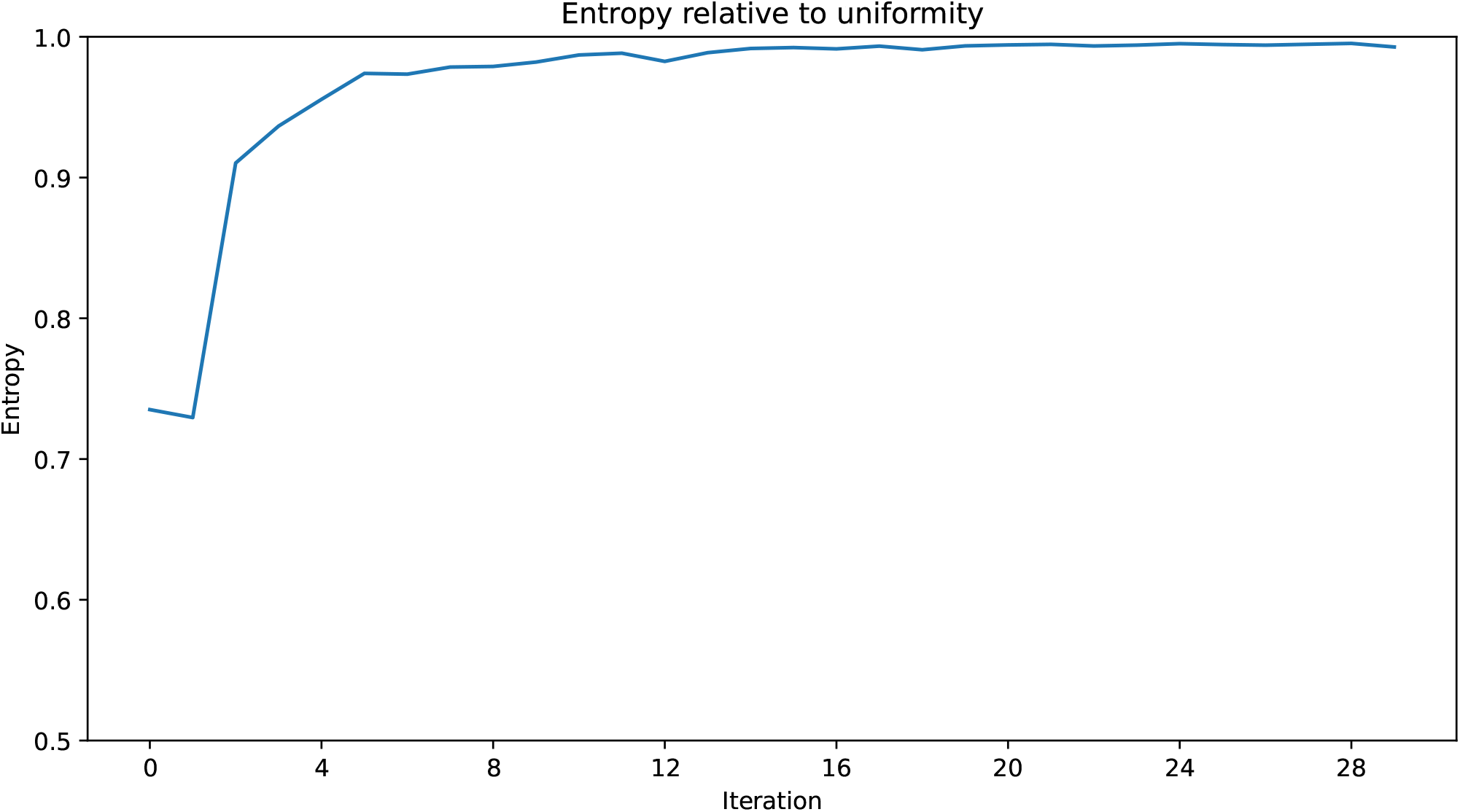
Entropy of the discriminator predictions relative to uniformity, for the isolation-with-migration model in each adversarial Monte Carlo iteration. When the discriminator predictions provide no additional information compared with the sampling distribution, the relative entropy tends towards 1. This measure was suggested by West (1993) as an indicator of convergence for an analogous iterative estimation algorithm, with the caveat that we cannot determine if the process has converged to the **correct** distribution.

**Figure S5:**
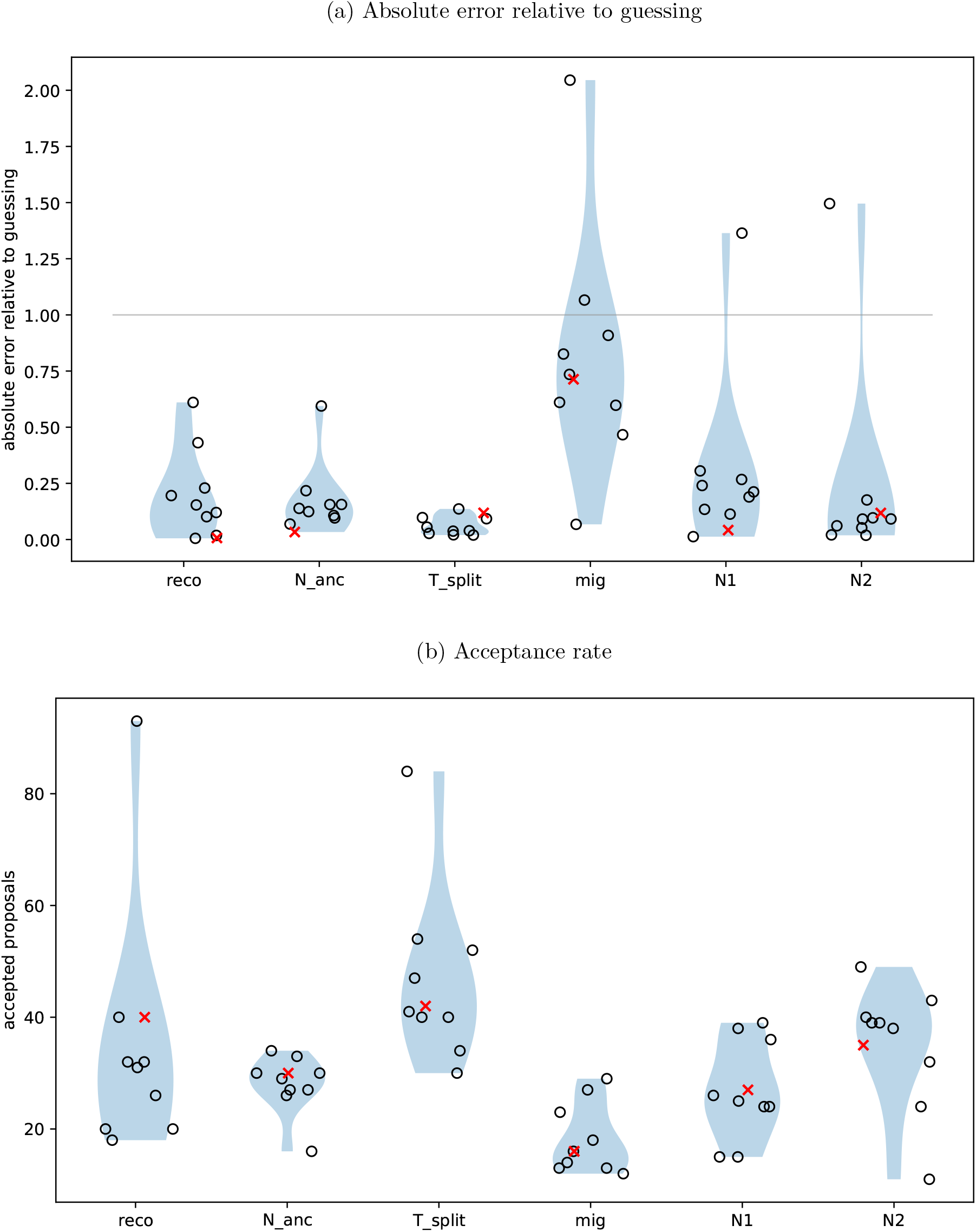
Results from 10 runs of `pg-gan` on the isolation-with-migration model. (a) The absolute error of each estimate, divided by the expected error from sampling uniformly at random—values lower than 1.0 are better than guessing. (b) The number of proposals that were accepted for each parameter, out of a total of 300 iterations. Red crosses indicate the `pg-gan` run with the lowest value for the generator loss.

